## Supplementary Information for "Context-aware deconvolution of cell-cell communication with Tensor-cell2cell"

**Supplementary Notes**

***Simulating 4D-communication tensors***

Tensor simulation consisted of six steps 1) generating the protein-protein interaction (PPI) network for ligand-receptor (LR) pairs, 2) generating the cell-cell (CC) network, 3) separately labeling LR pairs and single cells with a metadata group or category (to represent signaling pathways and cell types, respectively), 4) randomly associating a subset of LR pairs in the same signaling pathway with a subset of sender-receiver cell type pairs (generating a LR-CC combination), 5) assigning a context-dependent pattern of communication scores to LR-CC combinations, and 6) filling the tensor with communication scores that follow assigned communication patterns across contexts.

The LR pairs are generated as a random, unweighted, bipartite network; each of the two node types represents either a ligand or a receptor. This network retained a scale-free property using *StabEco*’s (v0.1) *BiGraph* function, with the power law exponent value set to 2 and the average degree value set to 3. We confirmed that the degree distribution does fit the power-law using a maximum likelihood estimation method from *igraph* (v0.8.3), and proceeded to remove all disconnected nodes from the network. We next generated an edge list of all cell-cell interactions, which is simply all pairwise permutations of cells; permutations retain directionality, allowing a distinction between sender and receiver cells. In distinct cell-cell interactions, autocrine interactions (self-loops) are allowed. LR pairs are uniformly binned to one of three metadata categories. The same process is repeated for cells, which are then further condensed from individual cells to their category to represent bulk rather than single-cell data; autocrine interactions at the single-cell resolution are considered homotypic interactions at the bulk resolution. Conceptually, these categories can be thought of as analogous to biological groupings such as signaling types (e.g., paracrine and endocrine) for LR pairs or cell-types for cells.

We simulate changes to intercellular conditions across twelve contexts. In order to populate the tensor, we randomly assign four communication patterns--expected communication scores that change as a function of condition--to distinct LR-CC combinations. An LR-CC combination” consists of one LR pair label (signaling pathway) and two cell labels (cell types), one for the sender cell and another for the receiver cell. In assigning patterns, signaling pathways or pairs of cell types may be individually reused in different communication patterns, but their combinations must be unique. For example, take two signaling pathways with the labels “X” and “Z”, and two cell types with the labels “A” and “B”. Accounting for directionality, cells can form the following interactions: “AA”, “AB”, “BB”, and “BA”. Next, any of the following LR-CC combinations may be assigned a pattern: “X-AA”, “X-AB”, “X-BB”, “X-BA”, “Z-AA”, “Z-AB”, “Z-BB”, “Z-BA”. If “X-AB” is assigned to a pattern, then tensor coordinates for LR-pairs categorized under the signaling pathway “X”, used by the sender cells categorized under cell type “A” and receiver cells categorized under cell type “B” will be filled with communication scores to reflect that pattern. LR-CC combinations not assigned to a pattern are considered the background. Biologically, the motivation for allowing redundancy in the LR-pair or CC-pair categories assigned to a pattern (e.g., “Z-AA” and “Z-AB”), is that the same groups of LR-pairs may dictate different types of interactions between different cell types (e.g., LR pairs under “Z” dictate a linear increase across contexts for “AA” cell interactions, but a pulse in “AB” cell interactions).

We then randomly initialized selected LR-CC combinations with one of four possible patterns (linear, oscillatory, pulsatile, and exponential) and their expected communication score in the first condition. Finally, we assign the expected communication score for each LR-CC combination in each proceeding context based on the respective pattern and starting communication score. All LR-CC combinations that were not chosen to have a pattern (the “background”) have an expected communication score of zero. Our final tensors have a size of 12 x 300 x 3 x 3 (contexts, LR pairs, sender cells and receiver cells, respectively).

### ***Adding noise to simulations***

To assess the robustness of our tensor factorization approach, we added noise to communication scores in each context and each pattern when filling the tensor. In the simulation, noise (represented as ***n***) is a parameter that can take on any value between 0 and 1. When ***n*** = 0, all communication scores are set to their expected value 𝞵. When ***n*** > 0, we draw from a uniform truncated normal distribution with the minimum and maximum values set to 0 and 1, respectively, and the mean seat to the expected value. The standard deviation 𝞼 is a function of noise ***n***. For 𝞵 > 0, 𝞼 = ***c·***𝞵***·n*** where ***c*** is a scaling factor which we set to 1.1. We represent this distribution as N~(𝜇 = 𝞵, σ =1.1𝞵***n****,* min=0, max=1). This causes the communication score dispersion to increase with noise (Supplementary Figure S14a).

At 𝞵 = 0, as in the background and in cases where LR-CC combinations assigned to a pattern reach an expected value of 0, we must adjust 𝞼 to be a function of a different value (notated 𝞵**_L_** instead of 𝞵); otherwise 𝞼 would always equal 0 independently of the value of noise. Specifically, 𝞵**_L_** represents the desired maximum average value of the communication scores at 𝞵 = 0, achieved when ***n*** = 1. We refer to 𝞵**_L_** as the maximum background noise. Thus, the distribution at 𝞵 = 0 is annotated as N~(𝜇= 0, σ =𝞵**_L_******n****,* , min=0, max=1). The entire distribution is then scaled by a factor **c’** to ensure that the average value of the distribution at ***n*** = 1, which we define as 𝞵**_B_**, is equal to 𝞵**_L_.** The need for scaling is apparent in its absence, or when setting **c’** = 1. In this case, 𝞵**_B_** is consistently less than 𝞵**_L_** at ***n*** = 1 (Supplementary Figure S14b). In order to identify an appropriate value for **c’**, we calculate 𝞵**_B_** at multiple values of 𝞵**_L_** ranging between 0 and 1, setting **c’** = 1. Next we fit a piecewise function to this curve, defined by the linear function 𝞵**_B_** = *m*𝞵**_L_** + *b* when 𝞵**_L_** < 𝞵* and the exponential function 𝞵**_B_** = $\left( \frac{\mu_{\boldsymbol{L}}-d}{c} \right)^{a}$when 𝞵**_L_** ≥ 𝞵*. We use *scipy’s* (v1.6.0) *curve_fit* function to estimate the parameters *m*, *b*, *d*, *c*, *a*, and 𝞵*. This fit allows us to predict the average value of the distribution N~(𝜇= 0, σ =𝞵**_L_******n****,* , min=0, max=1) when **c’** = 1 for a given 𝞵**_L_**. Annotating this prediction as 𝞵**^’^_B_**, we can then set **c’** = $\frac{\mu_{L}}{{\mu'}_{B}}$to achieve the desired behavior of 𝞵**_B_** = 𝞵**_L_** at ***n*** = 1 (Supplementary Figure S14c). Finally, since this scaling approach results in communication scores > 1, we set all drawn values greater than 1 to 1 (Supplementary Figure S14d). By limiting 𝞵**_L_** to values ≤ 0.25, we can avoid the bimodal distribution seen at 𝞵**_L_** ≥ 0.5. We can change 𝞵**_L_** for the background, which we refer to as the “maximum background score”, to any value between 0 and 0.25. For non-background contexts, when 𝞵 = 0, 𝞵**_L_** is set to the smallest non-zero 𝞵 for that specific LR-CC combination across all contexts. Allowing 𝞵**_L_** for the background to be changed in the simulation permits the independent assessment of noise added to the patterns only, or both the patterns and the background.

In the example tensor (Figure 1), we assigned a minimal baseline level of noise (***n*** = 0.01) to avoid any issues that may arise from sparsity due to assigning communication scores of 0 to the background. We scale communication scores of the background, i.e. LR-CC combinations that were not assigned to a pattern, in such a manner that the average value would be 0.05 when ***n*** = 1 (i.e., we set 𝞵**_L_** = 0.05).

### ***Accuracy of the tensor decomposition in assigning ligand-receptor importance***

To quantify the accuracy of the tensor decomposition in selecting important LR pairs for each factor, we used a Jaccard index. Here, we compared the LR loadings of the respective factor with the “ground truth” communication scores of LR pairs assigned to each pattern. To do so, we first binarized LR loadings in each factor by categorizing them into high or low loading equal-interval bins, under the assumption that the decomposition provides sufficient separation in loadings between those assigned to a pattern and those that are in the background. With these binarized scores, we can assume LR pairs in the high loading bin are influential for a particular factor. Finally, we computed the pairwise Jaccard index between LR pairs assigned to a given pattern from the simulation and “high-loading” LR pairs in each factor (Supplementary Table S1). We expected a high Jaccard value exclusively between an assigned pattern and the factor that has loadings in the context dimension that recapitulate that pattern.

We also used a Pearson correlation metric to compare LR loadings with communication scores assigned to each pattern. Since all patterns are assigned to unique LR-CC combinations, we can reduce the tensor to a matrix confined to the contexts and LR pairs for any given assigned pattern by selecting only the sender and receiving cells assigned to that pattern. We expect that LR pairs with high loadings have high variance in communication scores across contexts, since this variance reflects context-dependent changes. Thus, we calculated the pairwise Pearson correlation between the context-driven variance in communication scores across all LR pairs that were assigned to a given pattern and the LR loadings of each factor (Supplementary Table S2).

In the example simulated tensor (Figure 2e-f), the assigned communication scores for each pattern form a bimodal distribution. The corresponding loadings of each factor resulting from tensor factorization similarly demonstrated bimodal distributions. Pairwise comparisons between communication scores and factor loadings resulted in Jaccard indices of 1 for LR pairs assigned to a pattern and 0 for others; similar results were obtained for Pearson correlations.

### ***Assessing the robustness of the tensor decomposition to noise***

To assess the robustness of Tensor-cell2cell to noise, we simulated tensors, as described previously, at varying levels of noise; we iterated through noise values ranging between 0 to 1 and measured the decomposition error in each iteration. In each iteration, much like an elbow analysis, we decomposed the simulated tensor to ranks of 4, 5, and 6. Between the three decompositions, we retain the one that resulted in the smallest error, with the caveat that additional ranks should decrease error by at least 1.1 log-fold change (LFC).

We ran the analysis 1000 times to evaluate the effect of different random seeds for generating noise while holding the maximum background score constant. We repeat this process for multiple maximum background score values. Next, at each maximal background score, we fit a locally weighted smoothing (LOESS) curve to the error and noise outputs using statsmodels’ (v0.12.2) lowess function. This gave us predicted error measurements for each level of noise tested. To assess the amount of noise that needs to be added to the system to surpass a heuristic threshold of error--in this case 0.3--we then interpolated noise from the LOESS-predicted errors using scipy’s interp1d function.

### ***Tensor-cell2cell is robust to noise***

We generated 133,000 simulated tensors using the same parameters for generating the example tensor (Figure 1), but while also adding varying levels of noise to assess how the error of decomposition may be affected. To ensure that our assessment was independent of a specific interaction network, in each simulation, a new ligand-receptor PPI network was generated, and interaction patterns were assigned to new LR-CC combinations. Noise here was intended to represent non-biological variation that may arise in single-cell measurements due to a number of technical factors in RNA-sequencing, such as sample handling and library preparation. The addition of noise dampens the signal of each communication pattern in two ways: 1) by decreasing the change in communication score between conditions (resolution noise) and 2) by increasing the presence of interactions occurring by chance through randomly assigning higher communication scores to ligand-receptor and cell-cell pairs not assigned to a communication pattern (background noise).

When running Tensor-cell2cell on each of the simulated tensors, 85.1% of the decompositions resulted in a minimum error when the rank was set to equal the number of simulated patterns (r = 4) (Supplementary Figure S2a). Meanwhile, decompositions with ranks of 5 and 6 minimized error in 10.4% and 4.5% of cases, respectively. We assessed the level of noise required to surpass a heuristic error threshold of 0.3 at each level of maximum background noise (Supplementary Figure S2b). We found that this decomposition error was reached when adding a resolution noise between 0.26-0.37; the resolution noise needed to achieve an error of 0.3 monotonically increased with decreasing maximum background noise. Thus, Tensor-cell2cell is robust to both resolution and background noise. Taken together, these results indicate that Tensor-cell2cell is capable of handling noise in both the interacting and non-interacting cells when capturing various condition-dependent patterns.

***Tensor-cell2cell is fast and accurate***

We ran Tensor-cell2cell and CellChat on a single-cell transcriptome atlas of peripheral blood mononuclear cells (PBMCs) from COVID-19 patients with varying severity^1^ to measure the time and memory demands of each tool when performing the context-driven CCC analysis (Supplementary Figures S3a-d). Considering the number of samples in this dataset, processing time of CellChat scales more rapidly with the high number of pairwise comparisons. To control this, we varied the number of samples and performed this benchmarking in two scenarios: 1) by considering every sample individually as a context, wherein one can obtain sample-specific signatures that may coincide with others of the same severity (Supplementary Figures S3a-b), and 2) by considering every severity (control, mild/moderate and severe/critical) as contexts by aggregating cognate samples (Supplementary Figures S3c-d), which keeps the number of pairwise comparisons constant at three comparisons but at the expense of losing sample-specific information.

For a fair comparison of the tools, we set them to perform the analysis using exactly the same list of LR pairs, shared cell types across samples, as well as to run only one CPU core. Tensor-cell2cell performed better in all cases we tested exclusively running in the CPU (without the GPU scenario, Supplementary Figures S3a-d). The most favorable scenario led to an ~89-fold improvement in running time, which occurred when 60 samples were analyzed as individual contexts and CellChat comparisons were run under the “structural” method (computes a network topology dissimilarity^2^**).** However, CellChat performed only half as fast as Tensor-cell2cell when using the “functional” comparison (Supplementary Figure S3a), which is based on a Jaccard similarity. Other cases where CellChat was comparable to Tensor-cell2cell were when samples were aggregated into major contexts (Supplementary Figure S3c); however, this demanded substantial increases in memory usage (Supplementary Figure S3d). The “structural” method can handle cell types that are not present in all contexts, which may explain the inferior speed than the “functional” method. However, the comparison between the “structural” method and Tensor-cell2cell is pertinent since our tool can also handle cell types that are not present in all contexts, despite that the current algorithm is not optimized for this purpose as other tensor decomposition methods do, and both tools were set to run on just cell types that are in all samples. In addition, part of the speed-up of Tensor-cell2cell over CellChat could be due to the language used to build each tool (CellChat runs on R, while Tensor-cell2cell runs on Python).

Importantly, the improvement of Tensor-cell2cell is achieved even though it runs a brute-force elbow analysis (that is, by computing the error for every rank in a range of values). In this regard, this step can be either omitted (for example, when a desired number of factors is used; No elbow case in Supplementary Figures S3a-d) or optimized (for example, when a binary search is used), multiplying the speed-up we reported here by ~10-fold and ~3-fold, respectively. Remarkably, the memory usage of Tensor-cell2cell never surpassed 16GB in any of the tested scenarios, even when using 60 samples as individual contexts (Supplementary Figure S3b); meanwhile, CellChat surpassed 16GB when aggregating 12 samples (Supplementary Figure S3d) or using 24 samples as individual contexts (Supplementary Figure S3b). Again, this could be due to the programming language that each tool uses. Nevertheless, in terms of what each user would have to deal with, Tensor-cell2cell is more efficient in both time and memory, indicating it can more readily be run on multiple contexts simultaneously in a personal computer or laptop. Moreover, Tensor-cell2cell can run on a GPU when available, which can substantially improve the computational time of the analysis (up to 19- and 790-fold faster than the “functional” and “structural” methods of CellChat, respectively, when analyzing 60 PBMC samples; Supplementary Figure S3a-d).

We next evaluated the accuracy of Tensor-cell2cell and CellChat in classifying individual samples after predicting context-driven CCC (Supplementary Figures S3e-h). It is important to consider that the outputs coming from each tool are extremely different due to the scope of their analyses, so a direct comparison is not feasible. Hence, we instead used an intermediary approach that uses a classification model to evaluate how well each tool separates contexts given their outputs. In particular, we measured how well each tool separates samples by COVID-19 severity (Supplementary Figures S3e-f) and disease state (Supplementary Figures S3g-h). For this, we trained a classifier to predict severity (control, mild/moderate vs severe/critical) and a disease state (healthy vs COVID-19) in two different COVID-19 datasets, one containing PBMC samples^1^ and the other bronchoalveolar lavage fluid (BALF) samples^3^. We next measured their accuracy with the area under the receiver operating characteristic curve (AUC). Tensor-cell2cell outperformed CellChat when classifying PBMC samples by severity (Supplementary Figure S3e), and performed similarly when classifying samples by disease state (Supplementary Figure S3g). Moreover, Tensor-cell2cell performed better than CellChat in all classification tasks associated with BALF samples (Supplementary Figures S3f,h). Surprisingly, all methods performed better (highest AUC) when classifying BALF samples than when classifying PBMC samples, possibly due to a more evident severity-driven variation of the immune response in the infection site rather than in the periphery. Thus, these results show that Tensor-cell2cell can successfully find signatures of CCC that differentiate between contexts in a computationally efficient manner.

Importantly, the outputs and details offered by CellChat and Tensor-cell2cell differ. CellChat reports context-associated UMAP embeddings of signaling pathways, while Tensor-cell2cell outputs TCA embeddings for contexts, ligand-receptor pairs, and interconnected sender and receiver cells. By training classifiers that accept these differing outputs, in most cases, Tensor-cell2cell greatly outperformed CellChat (Supplementary Figures S3e,f,h). While in a few scenarios we observed qualitatively comparable performance (Supplementary Figure S3g, Tensor-cell2cell and the functional method of CellChat), Tensor-cell2cell always performed better quantitatively. Although context classification is a useful approach for comparison, this strategy cannot evaluate how well these methods infer CCC due to their distinct scopes and the vast differences in the tools’ outputs. In this regard, the outputs of Tensor-cell2cell seem valuable for identifying specific molecular targets and involved cells of a context-dependent module of communication, encompassing information beyond the scope of CellChat, which largely focuses on pair-wise, context-specific differences between signaling pathways. Nevertheless, to make a fair comparison of both methods, we did not use all of the high dimensionality that Tensor-cell2cell outputs offer, and used just the context dimension loadings.

***Tensor-cell2cell is robust to communication scoring inputs***

To further compare decomposition results between communication scoring methods, we modified the CorrIndex metric to score each dimension separately, and retain the score of the most dissimilar dimension. While this approach disregards the combinatorial effects of patterns across dimensions that are accounted for in decomposition, it is more stringent and brings focus specifically to dissimilarities between decomposition. With this modified metric, we found the average similarity score decreased from 0.82 to 0.64. NATMI and CellChat, in particular, were more dissimilar from the other methods (Supplementary Figure S5), which agrees with the fact that these two exhibited the lowest similarity score of 0.68 using the unmodified CorrIndex.

Next, we asked whether a specific tensor dimension is driving the observed dissimilarity. We ran the Corrindex comparing each method on each tensor dimension separately (Supplementary Figure S6). While the sample, sender cell, and receiver cell dimensions all exhibited high similarity (average scores of 0.94, 0.92, and 0.93, respectively), the ligand-receptor dimension had a substantially lower similarity. In fact, we found that our modified scoring approach consistently identified the ligand-receptor dimension as the most dissimilar dimension across all comparisons. This indicates that the ligand-receptor dimension drives dissimilarity in decomposition outputs. This makes sense since the ligand-receptor dimension reflects the raw score output by each distinct method, as well as previous benchmarking reports concluding that the scoring method used has a substantial impact on the predicted interactions^4^.

Given the high similarity of the other dimensions, as well as that of the unmodified CorrIndex, we conclude that Tensor-cell2cell is robust to differences at the ligand-receptor resolution, mitigating the propagation of differences within the ligand-receptor dimension to other dimensions and consistently identifying overarching communication patterns. We see that while the quantitative CorrIndex agrees visually with the qualitative consistency in factorization outputs (Supplementary Figure S14), we also see that there are some discrepancies at the cell-resolution, especially with NATMI. This is expected given differences in the ligand-receptor scores, which directly define the overall sender-receiver communication. The CorrIndex for sender- and receiver- dimensions is high because considering a single dimension independently of the other dimensions enables an optimal alignment that otherwise may not be possible; this pitfall is mitigated by the fact that the unmodified CorrIndex score is high. Interestingly, while CellChat is the most dissimilar at the ligand-receptor dimensions, NATMI is the most dissimilar for the other three dimensions, further reinforcing the importance of not only considering each dimension separately, but all dimensions in combination.

***Tensor-cell2cell detects traditional mechanisms associated with immune response to infections***

Factors computed by Tensor-cell2cell provide further insights about immune responses during SARS-CoV-2 infection. For example, factors 4, 6 and 7 are associated with lymphocytes (Figure 4a), particularly the communication of NK and T cells (factors 4 and 6), and B cells (factors 4 and 7). CD44 is predicted to be an important receptor on lymphocytes (factor 4 in Figure 4b), which consistently is key for cell migration^5^ and resolving lung inflammation^6,7^. Among the top-ranked signaling molecules in NK and T cells as senders (factor 6 in Figure 4b), we found CCL5 and GZMA (granzyme A), which are involved in immune cell activation and cytotoxic effector functions of CD8+ T cell and NK cells, events that are key in the control of viral infections^8–10^, as well as the interaction of PTPRC (CD45) with MRC1 (CD206), which regulates T-cell functionality^11^. Similarly, factor 5 seems associated with antigen-presenting cells such as mDC, macrophages and B cells as senders, especially through known interactions facilitating antigen presentation^12–15^ (e.g. CD99-CD99 as well as interactions between integrin ITGB2 and intercellular adhesion molecules ICAM1 and ICAM2). Therefore, our strategy can successfully detect meaningful biological processes and cell-cell interactions involved during disease progression.

***Tensor-cell2cell elucidates molecular mechanisms distinguishing moderate from severe COVID-19.***

Tensor-cell2cell recapitulated molecular findings such as the role of SEMA4D-PLXNB2 interaction promoting inflammation validated in another work^16^, interaction that Tensor-cell2cell revealed to be stronger in cases with more lung inflammation (severe cases) (Figure 4). Our method also associated macrophage CCC in severe cases with interactions between CCL2, CCL3, CCR1 and CCR5, that are main proinflammatory molecules in COVID-19 as observed in another work^17^, wherein their importance as potential therapeutic targets for diminishing COVID-19 severity was proposed^17^. Additionally, we identified novel CCC patterns and mechanisms regarding COVID-19 pathogenesis. For example, Grant *et al.* reported that CD206^hi^ alveolar macrophages participate in the immune response to SARS-CoV-2 infection^18^, but the underlying mechanisms mediating this response remain unclear. Factor 10 seems to extend the results presented by Grant *et al.* by showing that macrophage-expressed MRC1 (CD206) interacts with PTPRC (CD45) expressed by other cells (Figure 4a-b). Interestingly, the MRC1-PTPRC interaction mediating macrophage communication can promote immune tolerance^11^, which is consistent with factor 10 being associated with moderate cases, wherein anti-inflammatory macrophages (M2-like phenotype) seem to be characteristic. Remarkably, the source article of the BALF dataset^3^ reported that M2-like (anti-inflammatory) macrophages were present with higher frequency than M1-like (pro-inflammatory) macrophages in healthy and moderate COVID-19 patients, while M1-like macrophages were more frequent in severe COVID-19 patients^3,19^, supporting the results of Tensor-cell2cell. However, this work only detected differences in the cellular compositions detected by their markers, but did not provide a link with molecular mechanisms. Another example is a recent GWAS study that reported 13 significant loci associated with SARS-CoV-2 infection^20^, wherein ICAM1 popped up as an involved gene. Remarkably, Tensor-cell2cell assigned a high loading to the ITGB2-ICAM1 interaction in a communication pattern that seems to be associated with antigen presentation (factor 5, Figures 4a-b), providing further insights of its potential mechanism.

***ASD pathogenesis could be explained by factors 3 and 4***

A longstanding hypothesis for ASD pathogenesis is that in some subjects, neurons exhibit local hyperconnectivity, but deficits in longer range connections^21–23^. Neurons in cortical layers 2/3 and corticocortical projecting neurons in layers 5/6 are the main sender cells in factor 3 (Figure 5a and Supplementary Figure S9), suggesting that factor 3 may relate to local overconnectivity in ASD. Two key regulators of neurodevelopment Neuregulin 1 (NRG1) and Ephrin A5 (EFNA5), were the main ligands associated with sender cells. Erb-B2 Receptor Tyrosine Kinases (ERBB2-4) and ephrin receptors (EPHA3-5,7 and EPHB2), another class of receptor protein-tyrosine kinases, were the main receivers. Factor 3 receptors were broadly expressed across cell types. The neuregulin/ERB and ephrin systems play major roles in neuronal migration^24,25^ and axon guidance^26^, so reduced signaling in ASD may contribute to migration errors and local hyperconnectivity.

A second longstanding theory for ASD pathogenesis is that an imbalance between excitation and inhibition contributes to cortical dysfunction^27,28^. Inhibitory interneurons are the key receiver cell types in factor 4 (Figure 5a and Supplementary Figure S9), with parvalbumin interneurons (IN-PV), and SV2C-expressing interneurons (IN-SV2C) as the top ranked cells, suggesting that factor 4 could relate to excitation-inhibition imbalance in ASD. Pleiotrophin (PTN), Protein Tyrosine Phosphatase Receptor Type M (PTPRM), and Heparin Binding EGF Like Growth Factor (HBEGF) were the top senders in Factor 4. The main receivers were Anaplastic Lymphoma Receptor Tyrosine Kinase (ALK), PTPRM, and ERBB4. PTN is released by neural stem cells to promote the normal development of newborn neurons by binding to ALK, a receptor tyrosine kinase in the insulin receptor family^29^. The insulin/ insulin-like growth factor 1 (IGF-1) signaling pathway has previously been implicated as a potential therapeutic target for normalizing functional connectivity dysregulation in syndromic and idiopathic ASD^30–32^ and a pilot clinical trial studying the effects of IGF-1 as a pharmacotherapeutic for ASD showed promising results^33^. These results suggest that Tensor-cell2cell could have utility as a tool for identifying novel pharmacotherapeutic targets.

***Analysis of the BALF COVID-19 dataset with CellChat***

We ran the analysis that CellChat offers for pairwise comparisons of contexts using the BALF dataset on the same conditions as used with Tensor-cell2cell. To simplify the analysis and interpretation, samples with the same severity were aggregated (Supplementary Figures S10-13). From the joint manifold learning analysis on signaling pathways (Supplementary Figure S10a), CellChat groups functionally similar molecules, especially in the immune response they participate in (e.g. cell adhesion and cytokines). By inspecting the pairwise comparison on signaling pathways (Supplementary Figure S10b), pathways such as GDF, OCLN, SELL, LAMININ and SPP1 seem to increase as severity increases. They are associated with moderate cases in the comparison of healthy vs moderate COVID-19, and associated with severe cases in the comparison of moderate vs severe. This suggests a potential correlation between these specific pathways and COVID-19 severity. Particularly, growth differentiation factors (GDFs) are the ones that Tensor-cell2cell did not detect, and they correspond to stress-, infection-, and inflammation-induced cytokines that can suppress immune responses^34^, potentially explaining the severity association detected by CellChat. Similarly, Occludin interaction (OCLN-OCLN) was not detected by Tensor-cell2cell’s built-in scoring function, an interaction that corresponds to an airway tight junction that is one target of viral infections^35^. SELL, Laminins and SPP1 involving integrins were captured by Tensor-cell2cell (Figure 4). Although SPP1 specific interactions are not among the top-5 LR pairs, they are among the top-10 pairs; and other integrin interactions were detected by Tensor-cell2cell. Moreover, the most important changes of CCC detected by CellChat correspond to macrophage interactions (Supplementary Figure S11-13). Here, it is easily noticeable that autocrine interactions of macrophages, involving CCL2, CCL3, CCL7 and CCL8 with their respective receptors (CCR1, CCR2 and CCR5), are increased in more severe cases; result that is coherent with the factor 8 output by Tensor-cell2cell. Thus, CellChat can offer pathway-level details missed by Tensor-cell2cell, but without providing information about their specific cellular associations. Furthermore, CellChat misses other mechanisms that Tensor-cell2cell found (Figure 4). Nevertheless, CellChat is still a powerful tool that can detect some of the results we presented with Tensor-cell2cell, such as the role of SELL and LAMININ pathways, the association of MIF and healthy patients, and the role of macrophages in more severe cases, potentially explaining its good performance in the classification benchmarking (Supplementary Figures S3e-h). Importantly, these discrepancies may be mitigated by extending Tensor-cell2cell to use other communication scoring methods as input, e.g. CellChat’s communication probabilities (Figure 3a and Supplementary Figure S4).

Finally, a key conceptual limitation of CellChat and other communication scoring tools, as compared to Tensor-cell2cell, is the need for pairwise comparisons between contexts, which prevents the identification of communication patterns across contexts and results in an exponential increase in the number of results with increasing samples/contexts, even when aggregating samples by context groups (Supplementary Figures S10-13). Another limitation is the inability to consider all biological scales (ligand-receptor, cell-cell, and context) simultaneously, both of which reduce the interpretability of results.

**Supplementary Table S1. Jaccard index to evaluate the accuracy of Tensor-cell2cell on ranking ligand-receptor pairs**

| Pattern | Decomposition Factor | | | |
| --- | --- | --- | --- | --- |
|  | Factor 1 | Factor 2 | Factor 3 | Factor 4 |
| Pulse | 0.00 | 1.00 | 0.00 | 0.00 |
| Oscillation | 1.00 | 0.00 | 0.00 | 0.00 |
| Linear* | 0.00 | 0.00 | 1.00 | 1.00 |
| Exponential* | 0.00 | 0.00 | 1.00 | 1.00 |

* In this simulation, the same set of LR pairs was assigned to both the exponential and linear patterns (but in each case they were used by different sender-receiver cell pairs). Thus, when the exponential pattern has a high Jaccard index, the linear pattern is expected to have a high Jaccard index as well, and vice versa.

**Supplementary Table S2. Pearson correlation to evaluate the consistency between ground truth ligand-receptor pairs and LR loadings.**

| Pattern | Decomposition Factor | | | |
| --- | --- | --- | --- | --- |
|  | Factor 1 | Factor 2 | Factor 3 | Factor 4 |
| Pulse | -0.49 | 1.00 | -0.50 | -0.50 |
| Oscillation | 1.00 | -0.49 | -0.51 | -0.51 |
| Linear* | -0.51 | -0.50 | 1.00 | 1.00 |
| Exponential* | -0.51 | -0.50 | 1.00 | 1.00 |

* In this simulation, the same set of LR pairs was assigned to both the exponential and linear patterns (but in each case they were used by different sender-receiver cell pairs). Thus, when the exponential pattern has a high Pearson correlation, the linear pattern is expected to have a high Pearson correlation as well, and vice versa.

**Supplementary Table S3. Literature support for the top-ranked ligand-receptor interactions of the COVID-19 study case**

| **Factor** | **Ligand*** | **Receptor*** | **Reported role in immune response and/or COVID-19** | **Refs.** |
| --- | --- | --- | --- | --- |
| Factor 1 | CD99 | CD99 | CD99 is involved in the transendothelial migration of neutrophils and monocytes | ^36^ |
|  | MIF | CD74 & CD44 | During early infection in COVID-19, plasma MIF concentration is increased. Reported association between an early MIF response, organ malfunction and 28-day survival | ^37^ |
|  |  |  | MIF-CD74 promotes wound healing and recovery during lung injury, including injuries associated with viral infections | ^38,39^ |
|  |  |  | CD74 is involved in blocking the entry of coronaviruses through endosomes, including SARS-CoV-2 | ^40^ |
|  | MDK | NCL | Potential role of MDK in facilitating viral entry. Nucleolin (NCL) facilitates nucleocytoplasmic transport of MDK | ^41^ |
|  |  | ITGA4 & ITGB1 | During inflammation, MDK-integrin interactions facilitate neutrophil trafficking | ^42^ |
| Factor 2 | SEMA4D | PLXNB2 | SEMA4D-PLXNB2 interaction participates in epithelial wound repair | ^43^ |
|  |  |  | SEMA4D regulates allergic inflammation in lungs. Participates in neutrophil activation | ^44^ |
|  |  |  | SEMA4D-PLXNB2 interaction promotes inflammation and neurodegeneration in experimental autoimmune encephalomyelitis | ^16^ |
|  | SEMA4A | PLXNB2 | SEMA4A-PLXNB2 induces production of Th17 cytokines in CD4+ cells | ^45^ |
|  | - | SDC4 | Syndecan-4 (SDC4) contributes to the cell entry of SARS-CoV-2 and attenuates antiviral immunity | ^46^ |
| Factor 3 | SIGLEC 1 | SPN | Siglec-1 binds SPN (CD43) during viral infection and this interaction inhibits interferon gamma production in T cells | ^47^ |
|  | RETN | CAP1 | Resistin (RETN) cytokine signaling via its receptor, CAP1, activates multiple inflammatory signaling pathways in monocytes | ^48^ |
|  |  |  | RETN is a marker of neutrophil activation, which has an increased production in patients with critical COVID-19 | ^49^ |
|  | FN1 | - | Increased levels of FN1 in lungs of patients with lung fibrosis | ^50^ |
|  |  | ITGA4 & ITGB1 | α4β1 integrin (ITGA4 & ITGB1) is involved in the recruitment of leukocytes by activated endothelial cells, involving migration processes such as tethering, rolling, arrest and adhesion. Fibronectin is one of its ligands. | ^51^ |
|  |  | ITGA4 & ITGB7 | α4 integrins are a target of natalizumab, a drug for treating multiple sclerosis. This drug had a favorable outcome in a COVID-19 patient with multiple sclerosis | ^52^ |
| Factor 4 | COL9A2 | CD44 | CD44 is a cellular adhesion molecule and receptor for laminin and collagen, among other ligands. Increased expression of CD44 in bronchial samples from patients with critical COVID-19. Increased collagen in lungs during viral infection. Laminin is increased in serum of COVID-19 patients. CD44 participates in resolution of lung inflammation. | ^7,17,53–55^ |
|  | LAMB3 |  |  |  |
|  | LAMB2 |  |  |  |
|  | LGALS9 | CD44 | LGALS9-CD44 regulates the immune response. LGALS9 is expressed by induced regulatory T cells as a mediator of immune suppression, which can act on cells expressing CD44 | ^56^ |
| Factor 5 | ITGB2 | CD226 | When upregulated on CD8+ T cells, CD226 enhances their cytotoxic effector functions | ^57^ |
|  |  |  | CD226 is upregulated on CD8+ T cells of COVID-19 patients | ^58^ |
|  |  |  | CD226+ monocytes are increased in COVID-19 patients compared to healthy patients | ^59^ |
|  | CD86 | CTLA4 | CTLA4 is upregulated in CD8+ T cells present in lungs of COVID-19 patients. CD86 is upregulated in macrophages with M2-like phenotype present in BALF samples of patients with severe COVID-19 | ^60^ |
|  | ITGB2 | ICAM2 | Expression of ICAM2 is upregulated in lung epithelium infected by SARS-CoV2 | ^61^ |
|  |  | ICAM1 | High expression of ICAM1 in lung epithelium infected by SARS-CoV2 | ^61^ |
|  |  |  | Increased presence of ICAM1 in lungs of patients with COVID-19 | ^62^ |
|  |  |  | Increased presence of ICAM1 in serum of patients with COVID-19, and higher levels were observed in non-survivors than in survivors | ^63,64^ |
|  |  |  | Neutrophil cytotoxicity is mediated by the interaction between ITGB2 (CD18) and ICAM1 | ^65^ |
|  |  |  | ICAM1 is associated with COVID-19 by a GWAS study | ^20^ |
| Factor 6 | CCL5 | - | Increased level of CCL5 at the early stage of infection in patients with mild COVID-19 | ^66^ |
|  |  | CCR5 | CCR5 gene expression is increased in BALF samples of patients with COVID-19 | ^67^ |
|  |  |  | CCR5 Δ32 polymorphism is positively correlated with SARS-CoV2 infection and COVID-19 mortality rate | ^68^ |
|  |  |  | CCL5-CCR5 interaction induces migration of macrophages and NK cells | ^69^ |
|  |  | CCR1 | Protective role of CCR1 and CCR5 in a MA15-SARS-CoV mouse model infection, and in human DCs infected with SARS-CoV. | ^69^ |
|  |  |  | CCL5-CCR1 interaction induces migration of macrophages and NK cells | ^69^ |
|  | GZMA | F2R | GZMA is involved in the response of CD8+ T and NK cells to SARS-CoV-2, helping to distinguish healthy patients from COVID-19 patients. Protease-activated receptor 1 (F2R) has been identified as a potential target for therapeutic purposes in COVID-19 | ^9,70^ |
| Factor 7 | SELL | - | L-selectin (SELL) regulates neutrophil trafficking to sites of inflammation | ^71^ |
|  |  |  | High expression of SELL in a subpopulation of monocytes present in patients with COVID-19 | ^59^ |
|  |  | MADCAM1 | SELL-mediated lymphocyte rolling on MADCAM1 | ^72^ |
|  | CD22 | PTPRC | CD22 is involved in response of B cells to SARS-CoV-2. PTPRC (CD45) is an immune regulator associated with severity in COVID-19 | ^73,74^ |
| Factor 8 | CCL2 | CCR2 | Presence of CCL2 in plasma of patients with critical COVID-19 is higher than in plasma of patients with mild COVID-19 | ^75^ |
|  |  |  | Increased expression of CCL8 in postmortem lungs from patients with COVID-19 | ^76^ |
|  |  |  | CCR2 gene expression is increased in BALF samples of patients with COVID-19 | ^67^ |
|  |  |  | CCL2-CCR2 interaction induces migration of inflammatory monocytes | ^69^ |
|  | CCL3 | CCR5 | High expression of CCL3 in the respiratory tract of patients with COVID-19 | ^77^ |
|  |  |  | CCL3 gene expression is increased in BALF samples of patients with COVID-19 | ^67^ |
|  |  |  | CCL3-CCR5 interaction induces migration of macrophages and NK cells | ^69^ |
|  | CCL8 | - | Increased expression of CCL8 in postmortem lungs from patients with COVID-19 | ^76^ |
|  |  | CCR1 | CCL8-CCR1 interaction induces migration of T cells involved in Th2 response | ^69^ |
|  | CCL3L1 | CCR1 | High expression of CCL3L1 in BALF samples from patients with COVID-19 | ^78^ |
| Factor 10 | CD99 | PILRA | PILRA is an inhibitory receptor expressed on macrophages. CD99 is a ligand of this receptor. PILRA regulates the recruitment of neutrophils in inflammatory responses. A knock-out mouse model of PILRA had neutrophils with enhanced transmigration | ^79,80^ |
|  | LGALS9 | HAVCR2 | HAVCR2 (TIM-3), is involved in T cell exhaustion and tolerance. TIM-3 is involved in T cell exhaustion in a progressive way with the severity of COVID-19 | ^81,82^ |
|  |  |  | High expression of TIM-3 in CD8+ T cells of patients with severe COVID-19, its expression is higher in CD8+ T cells from BALF compared to CD8+ T cells from PBMCs | ^83^ |
|  | ANXA1 | - | ANXA1 orchestrates epithelial repair | ^84^ |
|  |  |  | A mimetic peptide of ANXA1 has been proposed as a potential treatment of severe COVID-19 | ^85^ |
|  |  | FPR1 | FPR1 promotes wound healing in the respiratory tract | ^86^ |
|  | MDK | LRP1 | LRP-1 is one of the main receptors of MDK. MDK promotes the recruitment of polymorphonuclear cells during an acute inflammatory response. A recent article suggested a potential role of MDK-LRP1 interaction in neutrophil infiltration and the neutrophil extracellular trap formation during COVID-19 | ^41,42^ |
|  | PTPRC | MRC1 | MRC1 is expressed on surfaces of macrophages. PTPRC (CD45) and MRC1 (CD206) interaction promotes immune tolerance. CD206^hi^ macrophages are involved in immune response to SARS-CoV-2 infection. CD45 is an immune regulator associated with severity in COVID-19 | ^11,18,74^ |

* Some ligand-receptor interactions are repeated in different factors, but in this table they are mentioned only in their first appearance. Repeated LR pairs in different factors can be seen in Figure 4b.

**Supplementary Table S4. Top-5 ligand-receptor pairs captured in each factor of the tensor decomposition of the ASD data set.**

| **Factor 1** | **Factor 2** | **Factor 3** | **Factor 4** | **Factor 5** | **Factor 6** |
| --- | --- | --- | --- | --- | --- |
| NEGR1 - NEGR1 | LAMA2 - SV2B | NRG1 - ERBB2&ERBB4 | PTN - ALK | NRG3 - ERBB4 | VEGFA - FLT1 |
| NRXN3 - NLGN1 | LAMA4 - SV2B | NRG1 - ERBB2&ERBB3 | PTPRM - PTPRM | NCAM1 - NCAM2 | PTPRM - PTPRM |
| NRXN1 - NLGN1 | LAMB2 - SV2B | NRG1 - ERBB3 | HBEGF - ERBB4 | CADM1 - CADM1 | VEGFB - FLT1 |
| CTN1 - NRCAM | LAMA1 - SV2B | NRG1 - ERBB4 | NRG3 - ERBB4 | NCAM1- NCAM1 | NRXN1 - NLGN2 |
| NCAM1 - NCAM1 | LAMA3 - SV2B | EFNA5 - EPHB2 | BTC - ERBB4 | CTN1 - NRCAM | PTN - NCL |

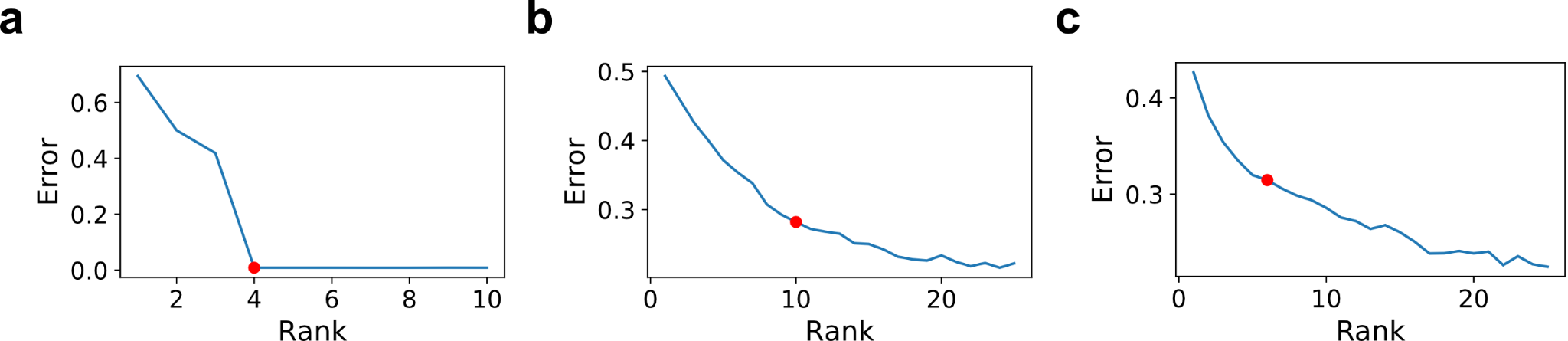

**Supplementary Figure S1. Rank selection for the tensor decomposition through an elbow analysis.** A selection of the rank employed for the tensor decomposition (i.e. number of factors) is obtained by an elbow analysis on a normalized error associated with the reconstruction of the original tensor (see the Methods section). This analysis was performed on the tensors built from (**a**) the simulated tensor of communication scores, (**b**) the BALF-COVID-19, and (**c**) PFC-ASD datasets. The red dot indicates the rank selected in each case for performing the decomposition analysis. The criteria here were selecting an error value close to 0.3 (or smaller if possible) and to the area with no substantial changes when increasing the rank used in the factorization (elbow), while also keeping the decomposition rank as low as possible.

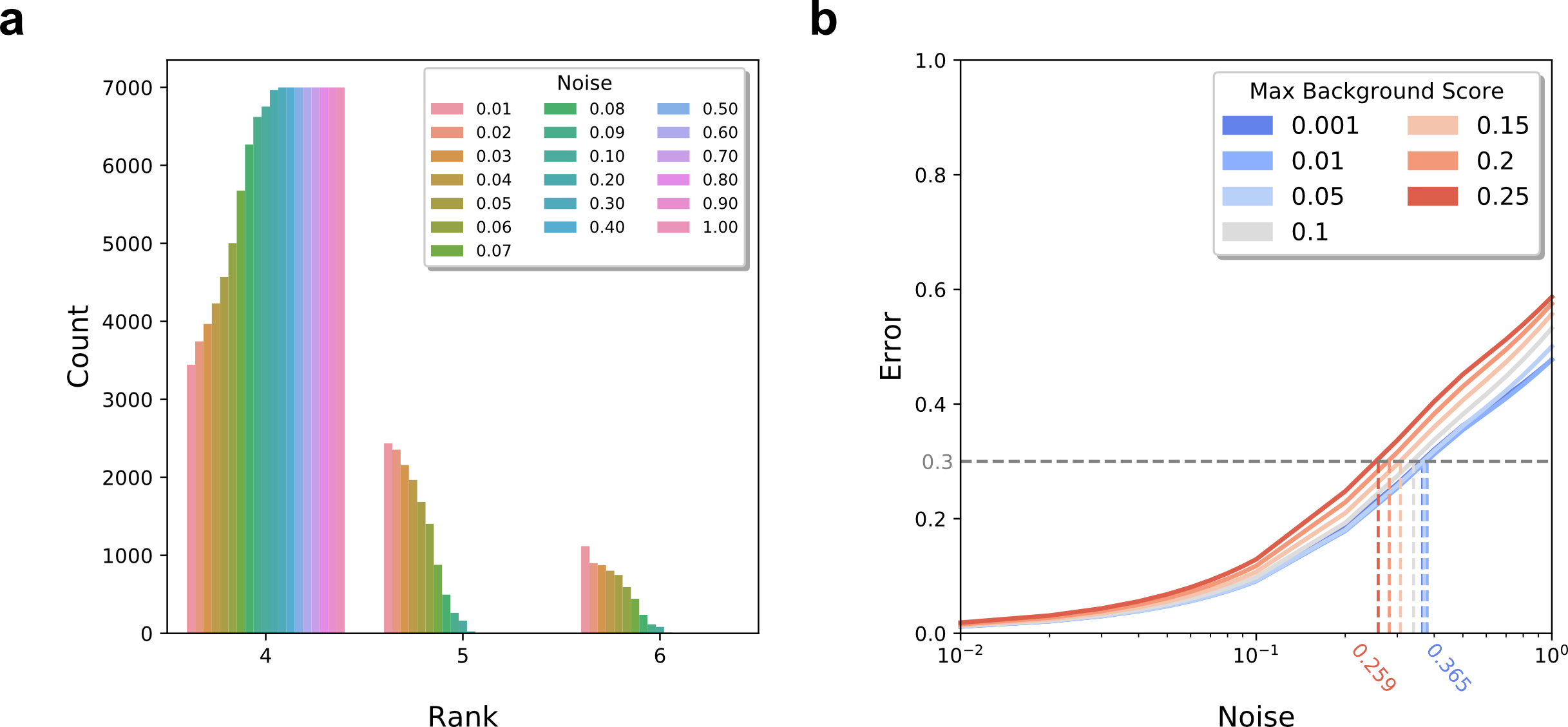

**Supplementary Figure S2. Tensor-cell2cell is robust to noise.** (**a**) Across all simulations, the total counts (y-axis) of each decomposition rank (x-axis) selected for error minimization. Counts within each rank are stratified by the level of noise (**b**) Locally weighted smoothing curves (LOESS, solid lines) visualizing the relationship between decomposition error (y-axis) and noise added to the communication scores during tensor simulation (x-axis). The maximum average communication score of the background in each simulation was limited to values ranging between 0.001 and 0.25 (legend). The amount of noise needed to achieve the heuristic error threshold of 0.3 (dashed lines) are interpolated from the LOESS predicted values for error. Interpolated noise values are displayed for maximum background noise values of 0.001 and 0.25.

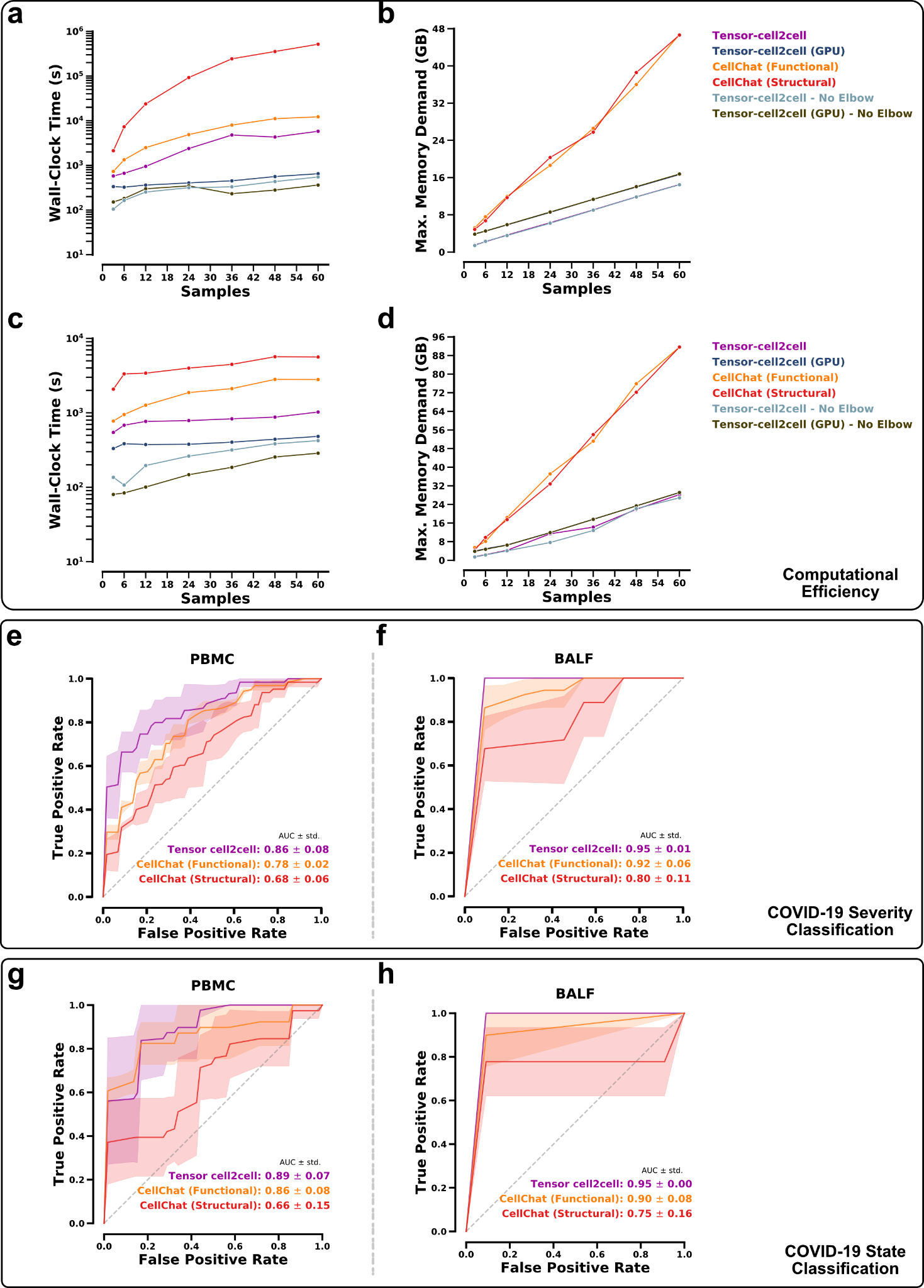

**Supplementary Figure S3. Benchmarking Tensor-cell2cell.** (**a**) Running time of Tensor-cell2cell and CellChat for analyzing CCC when the contexts correspond to individual patient samples. (**b**) Memory usage of each method to perform analyses in (a). (**c**) Running time of Tensor-cell2cell and CellChat for analyzing CCC when the contexts correspond to COVID-19 severity (individual samples were aggregated by severity: control, mild/moderate and severe/critical COVID-19). (**d**) Memory usage of each method to perform the analyses in (c). In (a) and (c), Tensor-cell2cell was benchmarked when running with or without a GPU, a feature unavailable in CellChat, and when considering or not the elbow analysis for selecting the number of factors. In addition, CellChat was benchmarked by using the two approaches it has for pairwise comparisons (functional and structural similarities, see Methods). (**e-h**) Receiver operating characteristic (ROC) curves of random forest models for classifying individual samples from the outputs of Tensor-cell2cell and the two CellChat approaches (functional and structural). These models predict specific severities of patients (control, mild/moderate or severe/critical) for (**e**) PBMC and (**f**) BALF samples, and disease state (healthy or COVID-19) for (**g**) PBMC and (**h**) BALF samples. For each classifier, the mean (solid line) ± standard deviation (transparent area) of the ROCs were computed from the 3-fold cross validations.

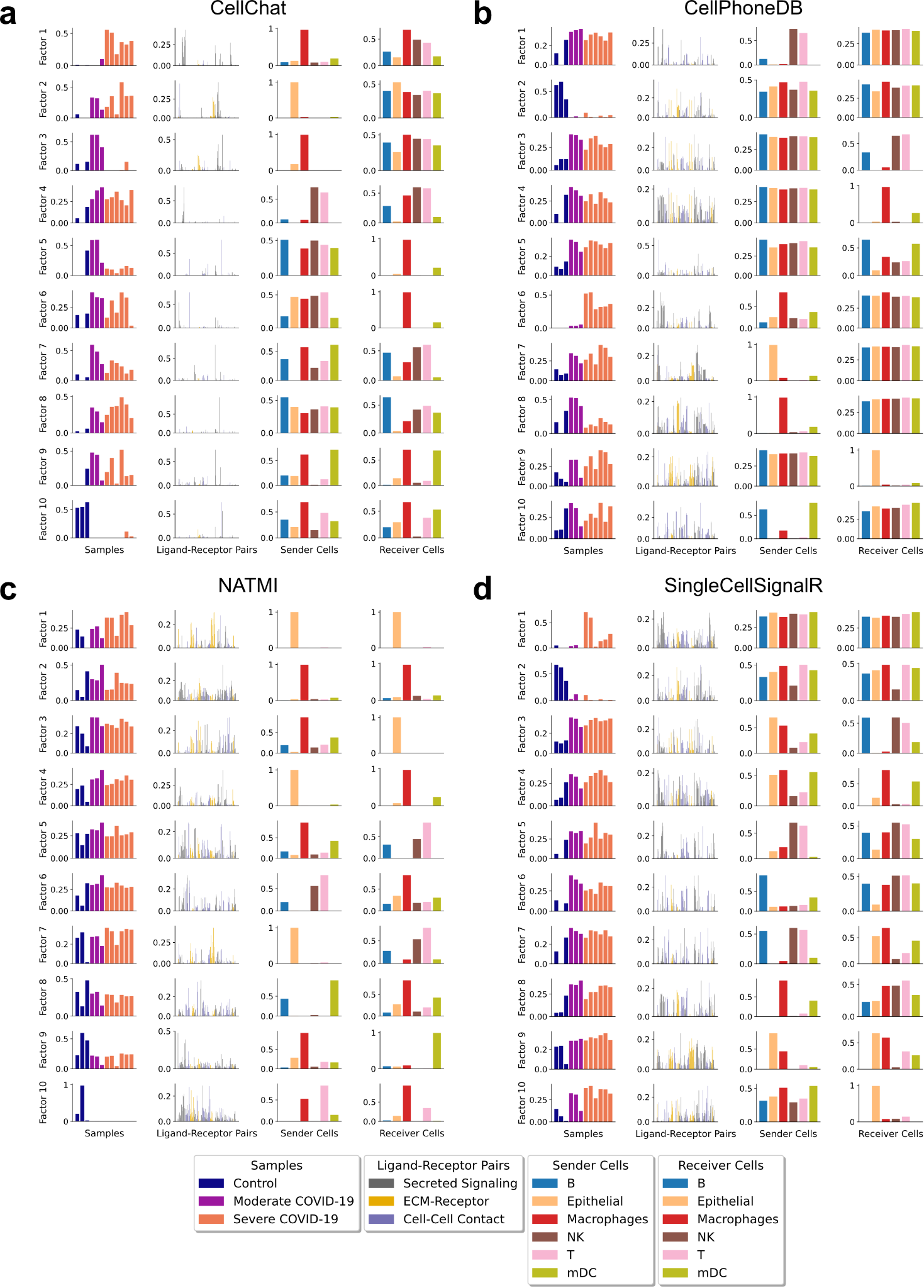

**Supplementary Figure S4. Tensor decompositions when using external tools to compute the communication scores.** Factors obtained after decomposing the 4D-Communication Tensors constructed from running external tools on each of the twelve BALF COVID-19 samples separately. The external tools used were (**a**) CellChat, (**b**) CellPhoneDB, (**c)** NATMI, and (**d**) SingleCellSignalR. For consistency, 10 factors (rank = 10) were selected for this analysis and 4D-Communication Tensors were subsetted to the same 176 ligand-receptor pairs, taken from CellChat’s database, and filtered for those present in all samples and scored across all tools prior to running decomposition.

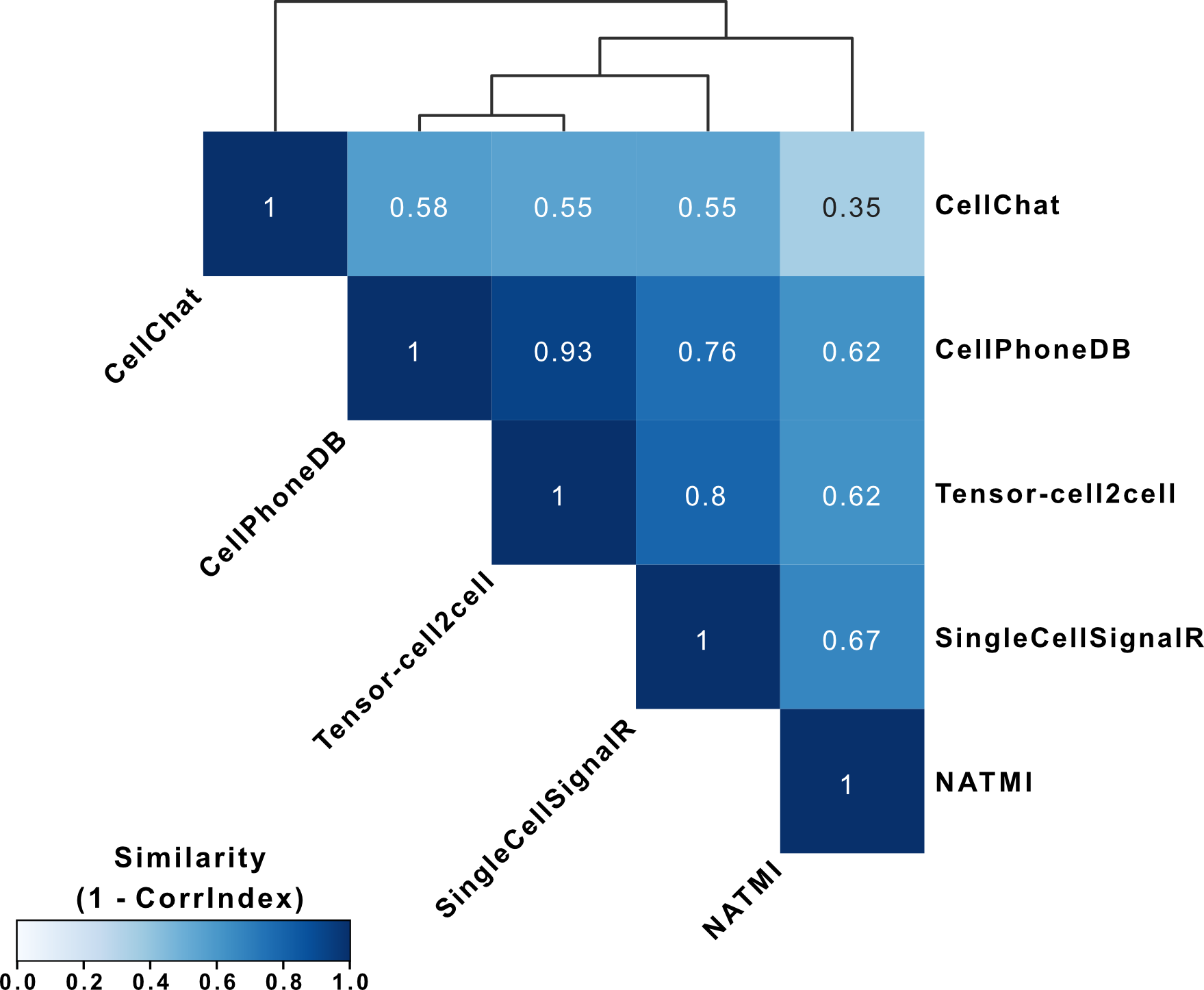

**Supplementary Figure S5. Conservative evaluation of decomposition similarity when using different communication scores.** Similarity (1 - CorrIndex) between tensor decompositions performed on the same 4D-communication tensor for a single-cell dataset of BALF in patients with varying severities, but with the communication scores computed from different tools for inferring cell-cell communication. Here, the CorrIndex score is modified for stringency by calculating it on each tensor dimension separately and subsequently selecting the maximal value (most dissimilar). Pairwise resultant similarities between all scoring methods are hierarchically clustered and visualized as a heatmap. Note that, since the ligand-receptor dimension was consistently the most dissimilar across all comparisons, this figure is the same as Supplementary Figure S6b.

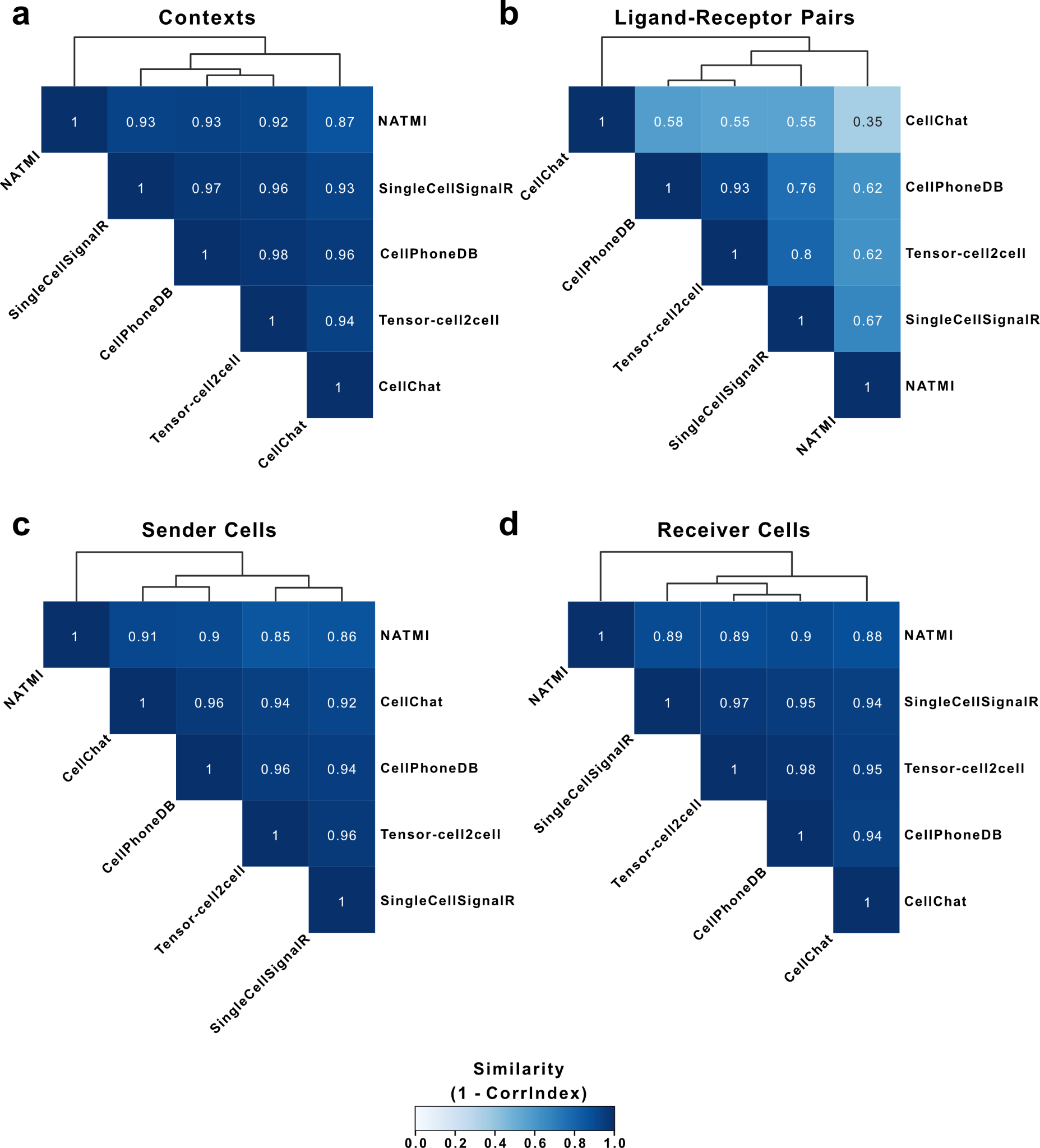

**Supplementary Figure S6. Evaluation of the main sources of dissimilarities when using different communication scores.** Similarity (1 - CorrIndex) between tensor decompositions performed on the same 4D-communication tensor for a single-cell dataset of BALF in patients with varying severities, but with the communication scores computed from different tools for inferring cell-cell communication. Here, the CorrIndex score is modified to consider each tensor dimension separately. Pairwise resultant similarities between all scoring methods for the (**a**) Context, (**b**) Ligand-Receptor Pair, (**c**) Sender Cell, (**d**) and Receiver Cell dimensions are hierarchically clustered and visualized as a heatmap.

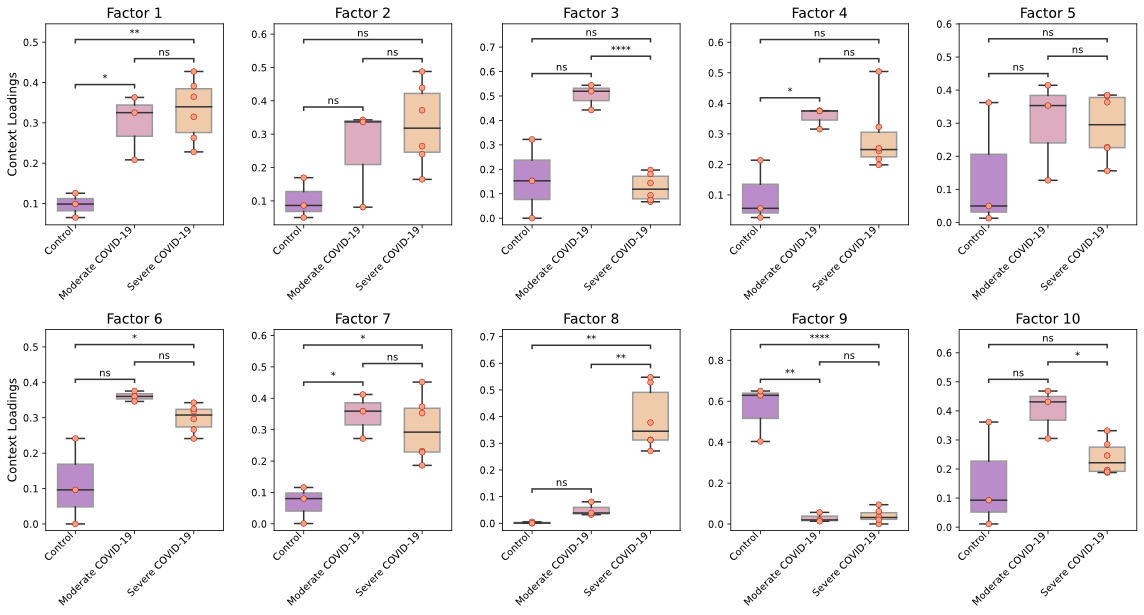

**Supplementary Figure S7. Boxplots of loadings for severities of COVID-19.** Boxplot representation for the different groups in the COVID-19 patients (healthy control, moderate disease and severe disease, as indicated in each x-axis). Each panel represents the sample loadings, grouped by disease condition, in each of the factors. Boxes represent the quartiles and whiskers show the rest of each distribution. A two-sided independent t-test was run to compare differences between the mean of the loadings of the groups, which was followed by a Bonferroni multiple test correction. For each pairwise comparison, the adjusted P-values are represented by: ns (P-value >= 0.05), * (0.01 < P-value < 0.05), ** (0.001 < P-value <= 0.01), *** (0.0001 < P-value <= 0.001) and **** (P-value <= 0.0001).

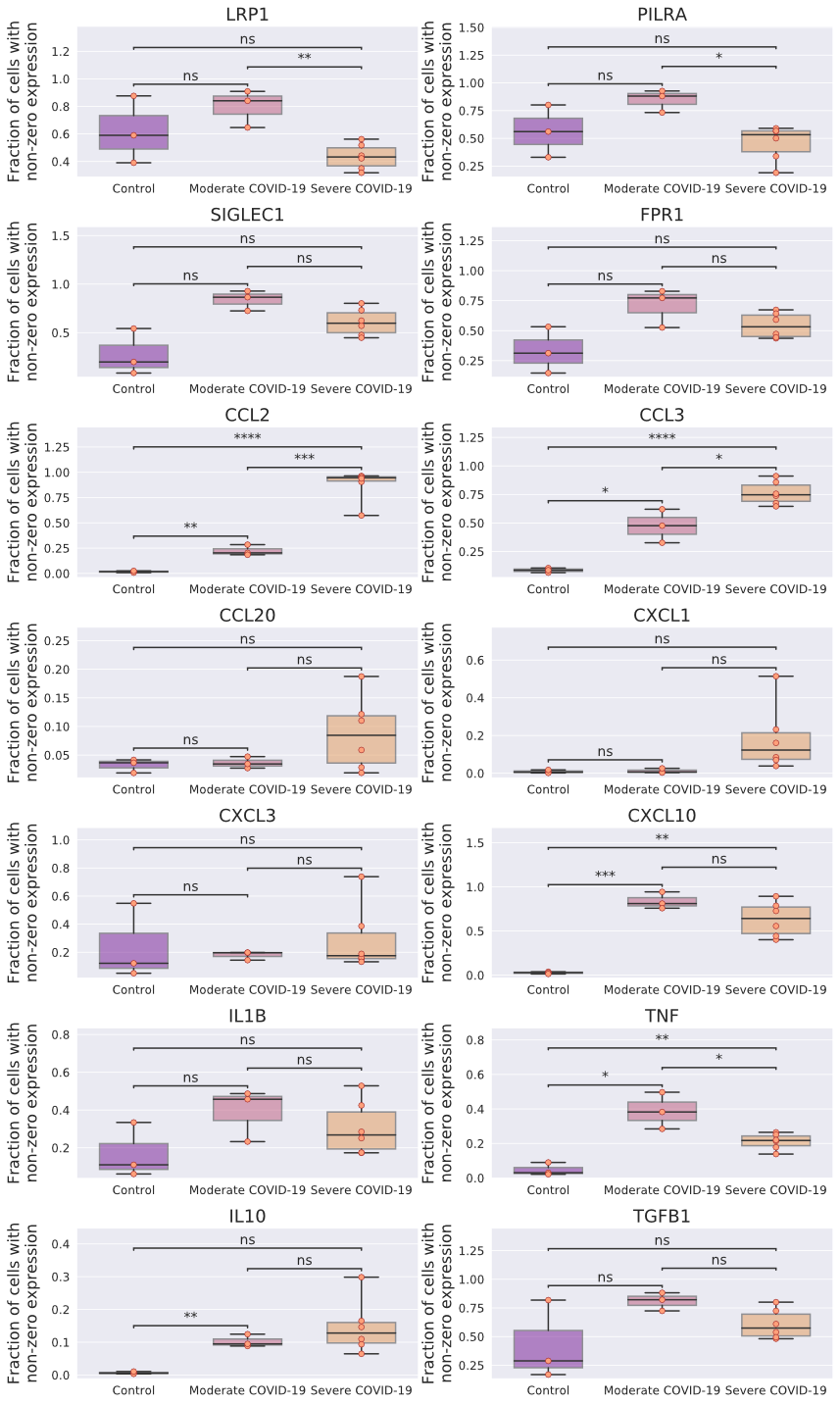

**Supplementary Figure S8. Gene expression associated with M1- and M2-like macrophages.** Gene expression of cytokines and/or receptors that are representative of M1- and M2-like phenotypes are shown for macrophages in each patient, grouped by COVID-19 severity (healthy group, moderate disease and severe disease, as indicated in each x-axis). Each subplot represents a different gene (as indicated in each title). In this case the gene expression values correspond to the fraction of cells with non-zero expression (y-axis) for the cluster of single cells annotated as macrophages. Boxes represent the quartiles and whiskers show the rest of each distribution. An independent t-test was run to compare differences between the mean of the loadings of the groups, which was followed by a Bonferroni multiple test correction. For each pairwise comparison, the adjusted P-values are represented by: ns (P-value >= 0.05), * (0.01 < P-value < 0.05), ** (0.001 < P-value <= 0.01), *** (0.0001 < P-value <= 0.001) and **** (P-value <= 0.0001).

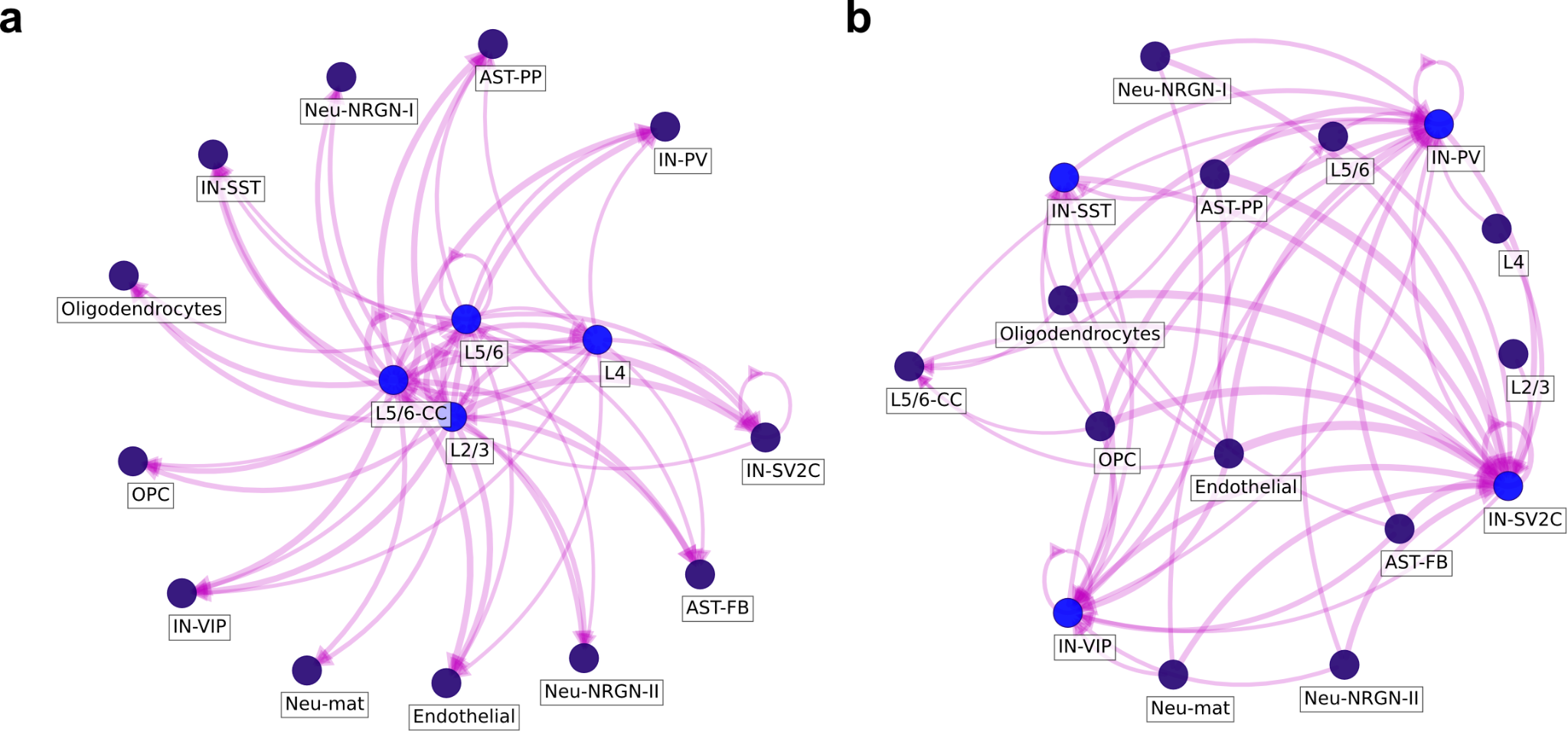

**Supplementary Figure S9. Factor-specific cell-cell communication networks that drive significant differences between ASD patients and controls.** Factor-specific cell-cell communication networks of (**a**) factor 3 and (**b**) factor 4 from the tensor decomposition of the ASD data set. These CCC networks representing the overall interactions between cells can be built for each of the factors by using the outer product between their respective sender-cell and receiver-cell normalized loadings (see ***Methods****)*. Resulting values represent edge weights. When building these networks, all cell types are connected to each other. To consider only biological meaningful interactions, we filter edges with a weight above a threshold of 0.075. In (**a**) and (**b**), nodes colored in blue represent those that are important as sender cells (factor 3) or receiver cells (factor 4). Edge widths are proportional to the edge weights.

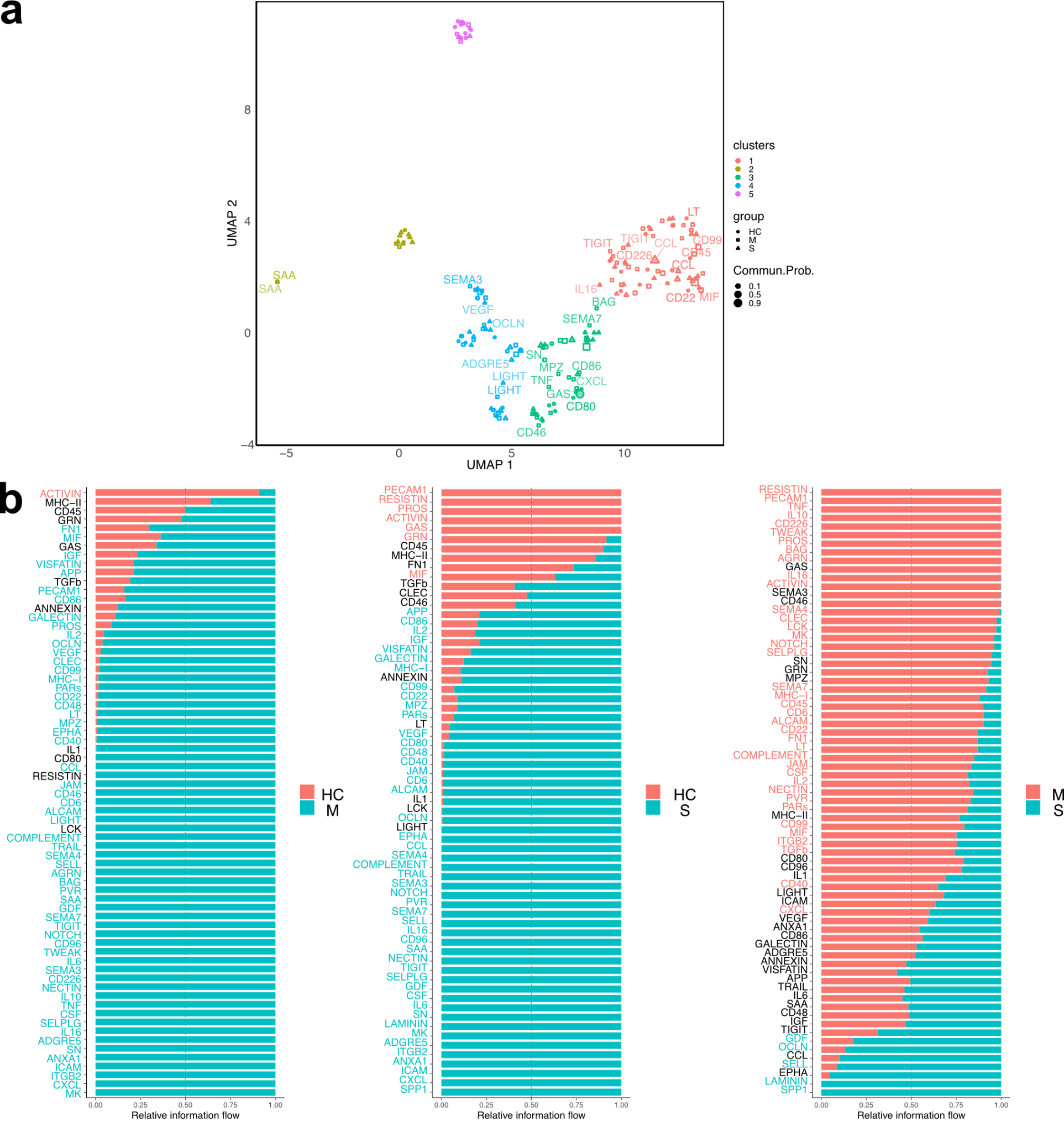

**Supplementary Figure S10. Downstream analyses of the BALF-COVID-19 dataset available to CellChat when considering multiple contexts.** For simplicity, sample expression matrices were aggregated by context (healthy control: *HC*, moderate COVID-19: *M*, and severe COVID-19: *S*) prior to running a CellChat functional analysis, as described in the ***Benchmarking of computational efficiency of tools*** section of the Methods. Analyses are conducted on the pairwise similarity between signaling pathways. (**a)** UMAP embeddings and clustering results computed from the pairwise similarity matrix between signaling pathways across all contexts. This manifold learning process summarizes multiple pairwise comparisons in an automated manner. **(b)** The ranked information flow (total communication probability between all pairs of cell groups) compared between each context for each signaling pathway.

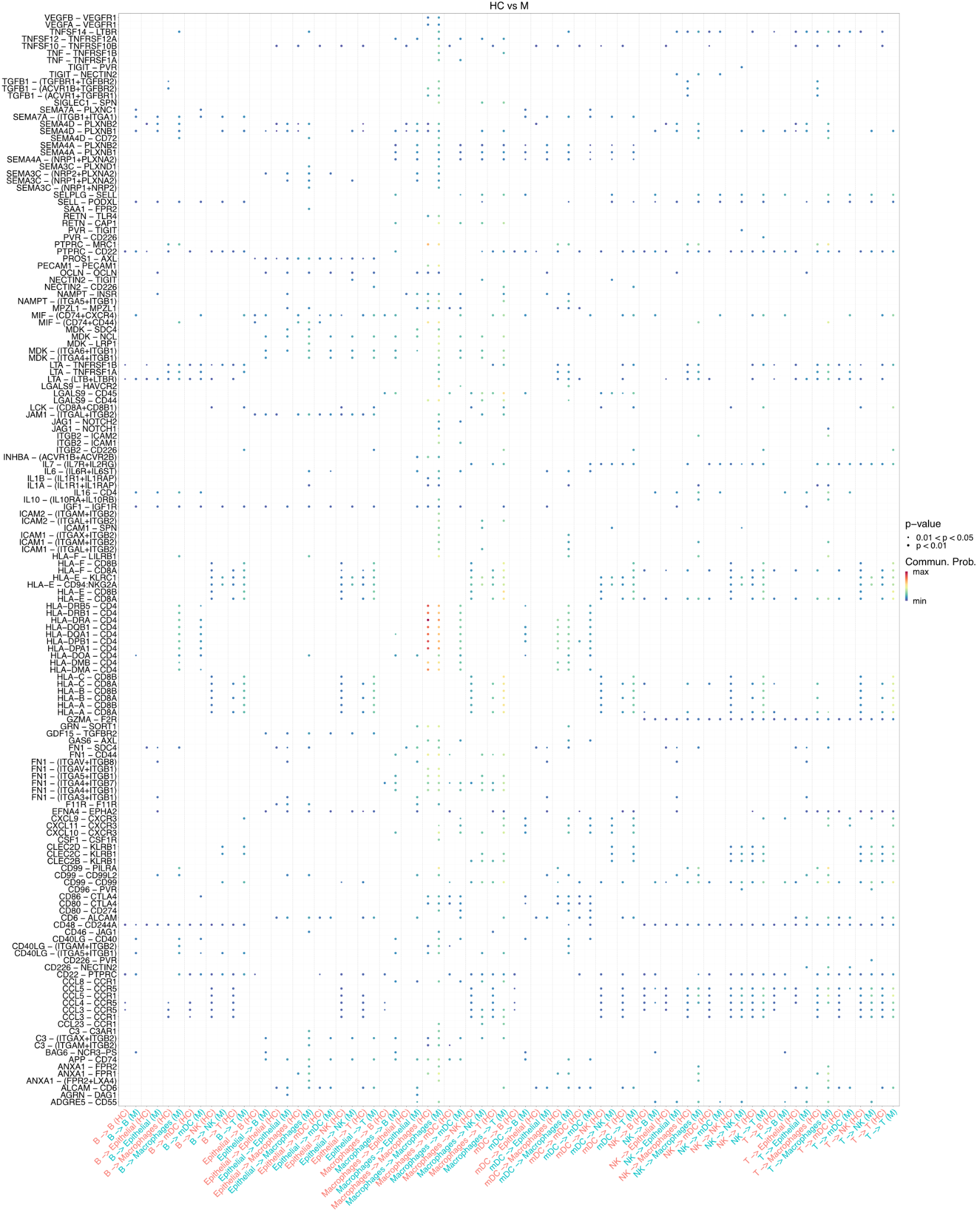

**Supplementary Figure S11. Comparison of communication probabilities for the BALF-COVID-19 dataset using CellChat.** Bubble plot generated with CellChat that shows the communication probabilities for a given sender-receiver cell pair and ligand-receptor pair, and that are directly compared between healthy (HC) and moderate (M) patients. Dot colors represent communication probabilities, and dot sizes are proportional to the P-values resulting from cell-type label permutation.

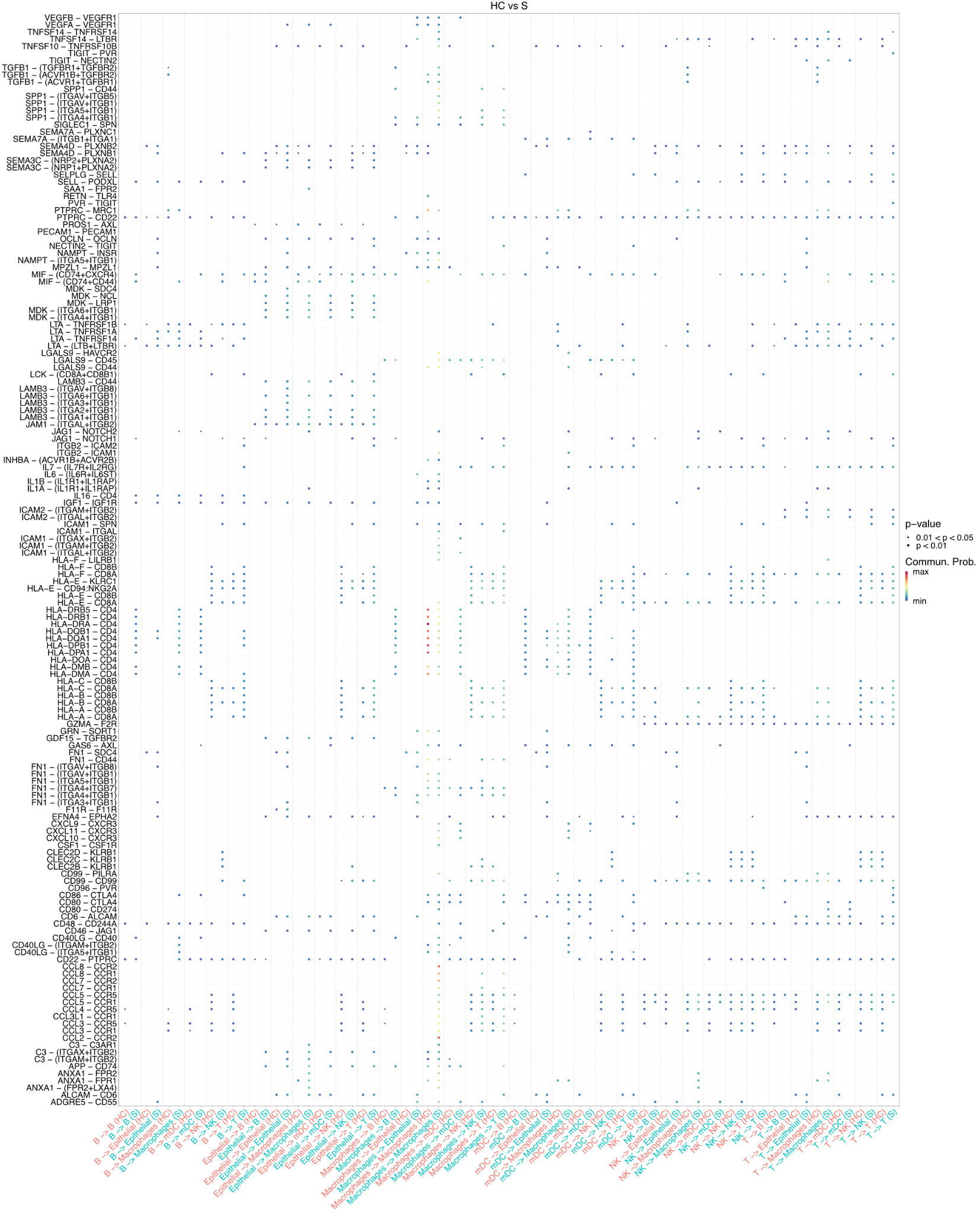

**Supplementary Figure S12. Comparison of communication probabilities for the BALF-COVID-19 dataset using CellChat.** Bubble plot generated with CellChat that shows the communication probabilities for a given sender-receiver cell pair and ligand-receptor pair, and that are directly compared between healthy (HC) and severe (S) patients. Dot colors represent communication probabilities, and dot sizes are proportional to the P-values resulting from cell-type label permutation.

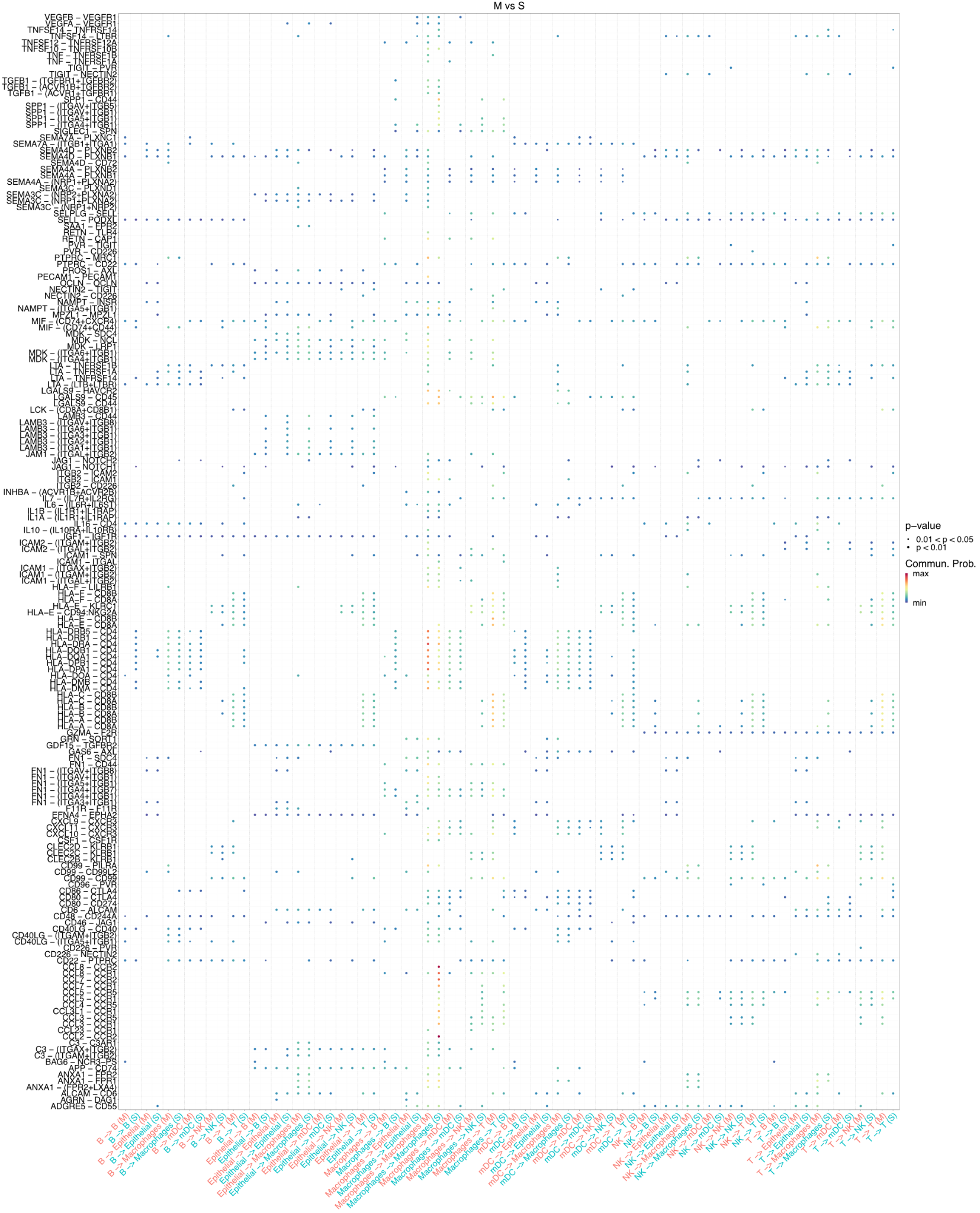

**Supplementary Figure S13. Comparison of communication probabilities for the BALF-COVID-19 dataset using CellChat.** Bubble plot generated with CellChat that shows the communication probabilities for a given sender-receiver cell pair and ligand-receptor pair, and that are directly compared between moderate (M) and severe (S) patients. Dot colors represent communication probabilities, and dot sizes are proportional to the P-values resulting from cell-type label permutation.

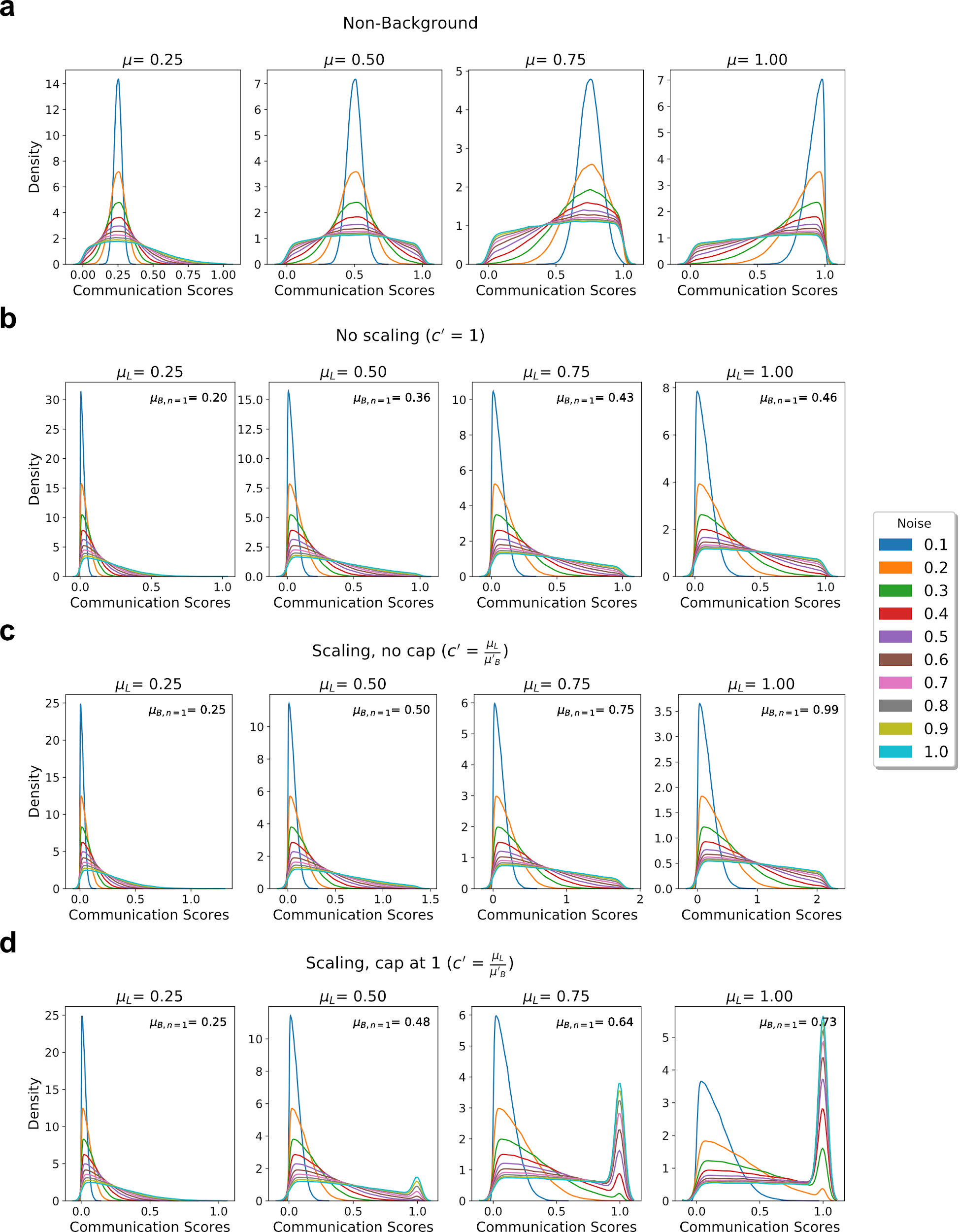

**Supplementary Figure S14. Noise increases dispersion of communication scores.** (**a**) From left to right, the distribution of communication scores at increasing levels of expected value across noise. (**b**) From left to right, the distribution of communication scores at increasing maximal average value when noise is 1, without scaling the distribution, for an expected value of 0. (**c**) From left to right, the distribution of communication scores at increasing maximal average value when noise is 1, scaling the distributions such that the average value of the distribution when noise is 1 equals the maximal average value, for an expected value of 0. Scores are not capped to have a maximum value of 1. (**d**) From left to right, the distribution of communication scores at increasing maximal average value when noise is 1, scaling the distributions such that the average value of the distribution when noise is 1 equals the maximal average value, for an expected value of 0. Scores are capped to have a maximum value of 1. Panels b-d are annotated with the average value of the distribution when noise is 1. 𝞵 represents the expected value of communication scores, 𝞵**_L_** is the desired maximum average value of the communication scores, 𝞵’**_B_** is a function of 𝞵**_L_** and **c’** is a scaling factor. All these parameters are described in more details in the Supplementary Notes (see ***Adding noise to simulations***).
